# Breeding for growth has resulted in increased tolerance to drought at the cost of genetic diversity in Scots pine

**DOI:** 10.1101/2023.09.14.557809

**Authors:** Francisco Gil-Muñoz, Haleh Hayatgheibi, Juha M Niemi, Lars Östlund, M Rosario García-Gil

## Abstract

Climate change possess a threat to forests and forestry. Drought has been identified as a one of the main issues due to its interaction with other biotic and abiotic stresses. Few studies have been done regarding breeding effect on the adaptability to climate change. After a common garden experiment with seedling families of Scots pine from northern Sweden, we have found differences in drought tolerance between families of breeding and natural origin. We performed a high throughput analysis-based phenotyping on both canopy and root traits. Root architecture traits might be related to drought tolerance and show moderate to high heritability values. The heritability of root architecture traits can be useful not only for drought but also for adaptability to other abiotic stresses. Analysis on architecture traits show that, especially on canopy-traits, families from breeding origins show less phenotypic variance than the ones from natural origins. The methodology employed to evaluate drought tolerance and plant architecture might be useful for future research and forest management focused on climate change adaptability.

## Introduction

Droughts, windstorms, and floods are becoming more frequent and severe as a result of climate change (Diffenbaugh, Scherer and Trapp, 2013; Seneviratne *et al*., 2014; IPCC, 2018; Spinoni *et al*., 2018), also in northern Europe (Venäläinen *et al*., 2020). Drought increases the susceptibility of forests to storm and wind damage (Gardiner *et al*., 2013; Csilléry *et al*., 2017), which are becoming the main causes of forestry losses in Europe (Senf *et al*., 2020; Patacca *et al*., 2023). Therefore, with the frequency of drought events increasing (Samaniego et al., 2018; Spinoni et al., 2018), integrating stress tolerance into forest silvicultural and breeding pratices is becoming increasingly important in order to preserve forests (Dale *et al*., 2001).

In the last decade, research on morphological drought response in plants has grown at an exponential rate. Studies conducted to investigate drought tolerance in various plant species have revealed that root characteristics such as main root diameter (Richards, Condon and Rebetzke, 2001), percentage of fine roots (Fitter, 2002), and root density and surface (Turner, Wright and Siddique, 2001; Vadez *et al*., 2013) are associated with improved performance during water stress (Wasaya *et al*., 2018), particularly in drought-prone and low-productivity areas (Kuijken *et al*., 2015).

For example, plants with deep and fine roots contribute to a better drought tolerance, as deeper roots would reach deeper water reserves (Pirtel *et al*., 2021) and fine roots are less prone to cavitation (Phillips *et al*., 2016). In conifers, needle morphology (Gebauer *et al*., 2015, 2019), root and branching architecture have been linked with drought tolerance (Moran et al., 2017; Baldi and La Porta, 2022). Other studies have even suggested needle lifespan during drought stress as a key factor for drought tolerance (Song *et al*., 2022).

Root shape adaption has received a lot of attention since the origin of plant science (Hales, 1727; Schimper, 1908; Cannon, 1911; Weaver, 1919; Cody, 1986; Wasaya et al., 2018). However, due to the complexity of the root traits and the technical challenges, investigating root characteristics remains difficult (Nielsen, Lynch and Weiss, 1997; Sharma and Carena, 2016). According to Lynch (1995) root system architecture is divided into four different aspects: morphology, topology, distribution, and architecture. Morphology refers to the surface features of a single root axis, topology to how roots are connected from a branching perspective, distribution to the presence of root positional gradients, and architecture to the spatial configuration of the root system or the explicit deployment of root axes.

Although attempts have been made to incorporate root characteristics as a selection criterion in tree improvement programs (Baldi and La Porta, 2022), the difficulties in assessing root properties has hampered them. In conifers, breeding for drought tolerance have primarily focused on individual traits such as root depth (Cregg and Zhang, 2001; Matías, González-Díaz and Jump, 2014; Kolb *et al*., 2016) or general plant growth (De La Mata, Merlo and Zas, 2014; Kerr *et al*., 2015), whereas efforts to improve drought tolerance based on root and/or canopy architecture have yet to be made. Similarly, the fact that root-related traits are affected by high genotype by environment interaction (GxE) (Orman-Ligeza *et al*., 2014) due to factors such as temperature (Nagel *et al*., 2009), water availability (Bengough *et al*., 2011), and nutrients (Paterson *et al*., 2006; Pearse *et al*., 2006; Yoneyama *et al*., 2007; López-Ráez *et al*., 2008), poses an additional challenge (Kuijken *et al*., 2015). However, the use of phenotyping platforms in different setups and environments has been proposed to improve heritability estimates by decreasing the level of GxE (Kuijken *et al*., 2015).

Scots pine (*Pinus sylvestris* L.) is one of the most economically important forest species in Fennoscandia, where a large proportion of commercial stands are regenerated with seeds collected from superior trees that have been optimized for growth. Previously, an unfavorable correlation between growth and solid wood characteristics in Scots pine (Hong, Fries and Wu, 2014) and other conifer species (e.g., (Chen *et al*., 2016)) has been documented. A decrease in wood density has been linked to higher vulnerability to cavitation and thus increased susceptibility to drought in forest tree species (Hentschel *et al*., 2014; Rosner, 2017). Larger trees have been also described as having greater water demand, which may result in structural overshoot, exacerbating forests’ susceptibility to drought (Liang *et al*., 2021). Furthermore, the effect of directional selection during breeding may have decreased the level of genetic diversity. Despite this association and the possible effect on the gene pool, it remains to be researched if those changes have affected forests resilience to drought, and even their capacity to adapt to future increase in drought intensity. Therefore, in this study, we have conducted a common garden experiment with Scots pine seedlings treated under controlled and drought greenhouse conditions as a proxy to investigate the effect of breeding on forests’ resilience and adaptability to drought.

## Material and Methods

### Plant material

In the natural Scots pine stands, pine cones were collected from twenty trees, aged 30 to 250 year-old, at three different regions in Northern Sweden (ie., 120 families in total). Jokkmokk natural stand (Karatj-Råvvåive 66º41’14.2”N 18º56’37.4”E), Arjeplog natural stand (66º18’15.8”N 18º21’6.5”E), and Jämtland natural stand (Källberget-Storberget, 63º23’52.1”N 15º28’0.6”E). The first two locations are located further north than the last one. The seeds collected from the natural stands always represent open-pollinated families.

The improved genetic material consisted of seeds from full-sib and half-sib families obtained by the Swedish forestry research institute (Skogforsk) after controlled crossing of superior trees originating from the three same natural stands mentioned above. The total number of progenies involved in this study is included in Supplementary Table 1.

### Greenhouse conditions and drought stress

Seeds were planted in a mixture of 2:1 of garden soil (Planteringsjord, Plantagen, Sweden) and sand (Sandlådesand, Boke, Sweden) in volume. Seeds were germinated in AirBlock 121 (BCC AB, Sweden) trays of 54cc volume per cell. Trays were placed above a water flow restriction matrix of floral foam blocks (Oasis, USA) to simulate a water table as described in Marchin et al. (Marchin *et al*., 2020)). A total height of 3 and 21 cm between the bottom of the tray and the water level were used as control and drought conditions, respectively. In total, 24 seeds from each family were sown in a completely randomized five-block design. Four trays per block, two for drought and two for control. Between the foam and the trays, we placed 1 cm of sand to ensure contact between the trays and the matrix. Seeds were sown in May 2023 under greenhouse conditions, where temperature was maintained between 25 and 35°C and a 16:8h light:darkness photoperiod with Fiona Lightning (FL300) light from Senmatic in sunlight mode. The outer most cells of the trays were filled with soil but no plants were grown to avoid the border effect. Before the drought experiment started, all the seedlings were fertilized once a week by top watering for 35 days (HORTO LIQUID Rika S 7-1-5 at 10 ml/l concentration). The drought and control treatments lasted 45 days, during which time the plants were not irrigated nor fertilized from above. Water availability was measured during the entire period using a tensiometer (Supplementary Figure 1)

### Germination rate

Germination greatly varied between families, ranging between 0% and 100% depending on the family. In average 30% of families had germinated seeds. The average time between sowing and germination time was 16 days. More than 95% of the plants germinated between 12 and 23 days after sowing. Families with less than three plants in both control and drought were discarded from further analysis. No families were obtained from the natural stand in Jämtland due to insufficient germination rate. When the drought treatment began, there were 224 seedlings under control and 350 under drought that met the requirements indicated.

### Image acquisition

At the end of the experiment, plants were collected carefully removing the soil and placing them in a self-built acrylic cuvette with 1 cm of water to ensure the full expansion of roots (i.e., roots showed their natural shape) without tension similarly as described by York (2023). The cuvette was placed over a scanner (model) and each of the plants were scanned at 600 dpi (Supplementary Figure 2). A blue background was applied to facilitate image segmentation. Images were cropped during preprocessing and each of the scan images were divided between root and canopy. For image segmentation, a machine-learning approach using Ilastik software (Berg *et al*., 2019) was used to assess both canopy and root. Probability maps were used as input in the root analysis software that was used both for roots and canopy traits. The software used for root analysis was Rhizovision Explorer (Seethepalli *et al*., 2021). Settings for analysis were: Whole Root, Image Thresholding Level 120, keep largest component, Edge smoothing threshold 2 and Root pruning threshold 1.

### Root and canopy parameters

#### After image acquisition, the following parameters were scored for further analysis

Root parameters (as described in (Seethepalli *et al*., 2021):

Median Number of Roots, Number of Root Tips, Total Root Length, Root Depth, Root Width to Depth Ratio, Root Network Area, Root Convex Area, Root Solidity, Lower Root Area, Root Average Diameter, Root Perimeter, Root Volume, Root Surface Area, No. of root Holes, Average Hole Size, Average Root Orientation, Root Shallow Angle Frequency, Root Medium Angle Frequency, Root Steep Angle Frequency.

Furthermore, we calculated the following parameters for the root system: Prop Surface Area Diameter Range 1 (Proportion of root surface comprised by roots with diameter under 0.25mm), Prop Surface Area Diameter Range 2 (Proportion of root surface comprised by roots with diameter between 0.25 and 0.5 mm), Prop Surface Area Diameter Range 3 (Proportion of root surface comprised by roots with diameter between 0.5 and 1 mm) and Prop Surface Area Diameter Range 4 (Proportion of root surface comprised by roots with diameter between 1 and 1.5 mm).

Canopy parameters were calculated with a similar pipeline as the root parameters and the following were used: Canopy Height, Maximum Canopy Width, Canopy Width to Depth Ratio, Canopy Convex Area, Canopy Solidity, Needle Average Diameter, Needle Median Diameter, Canopy Perimeter, Canopy Volume, Average Needle Orientation and Canopy Surface Area

Total volume was just the addition of root volume and canopy volume. Ratios between canopy and root volume and canopy and root surface were also calculated.

### Statistical analysis

Statistical analysis was performed using RStudio (RStudio Team, 2021; R Core Team, 2022) v. with the following packages: dplyr, ggplot2, tidyverse, corrplot, stringr, factoextra. To calculate the drought tolerance index, a linear regression was used for the total volume in drought and control conditions for each family. The index is calculated based on the studentized residuals of a mathematical “second regression” relationship between the trait under control and drought conditions (Bidinger *et al*., 1982):

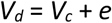

Where *V*_*d*_ is the family average of the total volume of the seedling in drought conditions, *V*_*c*_ is the family average of total volume in control conditions and *e* is the random residual effect. The deviation of each family from the model was used as tolerance index (underperformance and overperformance in drought conditions). Effect of the origin of the seeds was calculating adding an origin effect into the linear equation with interaction.

For testing the effect of family, origin and location, analysis was done using a General Linear Models (GLM). Gaussian, Poisson or gamma distributions were used accordingly to the distribution of the measured traits.

#### We fitted the data to the following statistical models

##### Family model (ANOVA F-test)

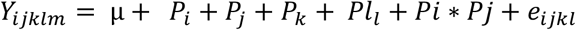

Where *Y*_*ijklm*_ is the trait phenotype on the *m*th seedling, *μ* is the overall mean of the response, *Pi* is the fixed effect of the *i*th family, *Pj* is the fixed effect of the *j*th treatment, *Pk* is the fixed effect of the *k*th block, *Pll* is the random effect of the age of the plant after germination (Days After Germination), *Pi* * *Pj* is the interaction effect between the *i*th family and the *j*th treatment and *e*_*ijklm*_ is the random residual effect.

##### Origin model (GLM)

In the case of TD, which consists of a single data point per tree, the model is *Y*_*ijklmn*_= μ + *P*_*i*_ + *P*_*j*_+ *P*_*k*_+ *P*_*l*_ + *P*_*lm*_ + *e*_*ijklm*_, Where *Y*_*ijklmn*_ is the trait phenotype on the *n*th seedling, *μ* is the overall mean of the response, *Pi* is the fixed effect of the *i*th origin, *Pj* is the fixed effect of the *j*th treatment, P*k* is the fixed effect of the *k*th region, *Pl* is the fixed effect of the *l*th block, *Plm* is the random effect of the age of the plant after germination (Days After Germination) and *e*_*ijklm*_ is the random residual effect.

For the exploratory analysis, a Principal Component Analysis (PCA) was done for all the traits both in control and drought for each family and another PCA was made for root traits at individual level of each seedling.

### Heritability estimates

Variance components and heritabilities for the investigated traits were estimated through four categories, comprised of: 1) progenies in control condition (seven full-sib and 26 half-sib families), 2) progenies in drought condition (seven full-sib and 30 half-sib families), 3) naturals in control condition (26 half-sib families), 4) naturals in drought condition (35 half-sib families).

A mixed-linear model approach, implemented in the ASReml4 statistical software package (Gilmour *et al*., 2015) was used following the model below:

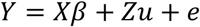

Where *Y* is the vector of observations; β is the vector of fixed effects (i.e., overall mean, block, days after germination); *u* is the vector of the random additive genetic effect of an individual tree; and *e* is the vector of random residual effect. *X* and *Z* are the incidence matrices relating the observations in *Y* to *β* and *u*, respectively. All random effects were assumed to be independently and normally distributed with the expected mean of zero where *var* (*u*) = *Aσ*^2^ and *var* (*e*) = *e*^2^ and *A* is the pedigree-based numerator relationship matrix.

Individual-tree narrow-sense heritability estimates (*h*^2^) were calculated as follows:

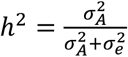

Where 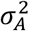 and 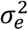 are the additive genetic and error variance components, respectively. The standard errors for variance components and genetic parameters were estimated by using Taylor series approximation.

## Results

### Drought stress effect on root and canopy

All the measured parameters were affected by drought except Canopy Solidity, Needle Average Diameter, Root Convex Area and Root Medium Angle according to the linear model (Supplementary Table 2). Canopy related variables, Width to Depth Ratio and Needle Angle, increased in response to drought treatment. Meanwhile, the rest of the canopy variables showed inferior values in drought. Under drought conditions, the root system tended to be smaller, both in surface and volume, and less compact but deeper. Root system had less tips, but the roots tended to be more horizontally inserted in comparison to the control. Drought also resulted in a higher proportion of roots with smaller sections than in control (Supplementary table X). Both Canopy to Root Surface and Canopy to Root Volume decreased in drought conditions because of a bigger decrease in canopy surface and volume as compared to the decrease in root under drought conditions. As an example, an average response of the canopy and root system to drought is represented in Figure 1.

**Figure 1.**
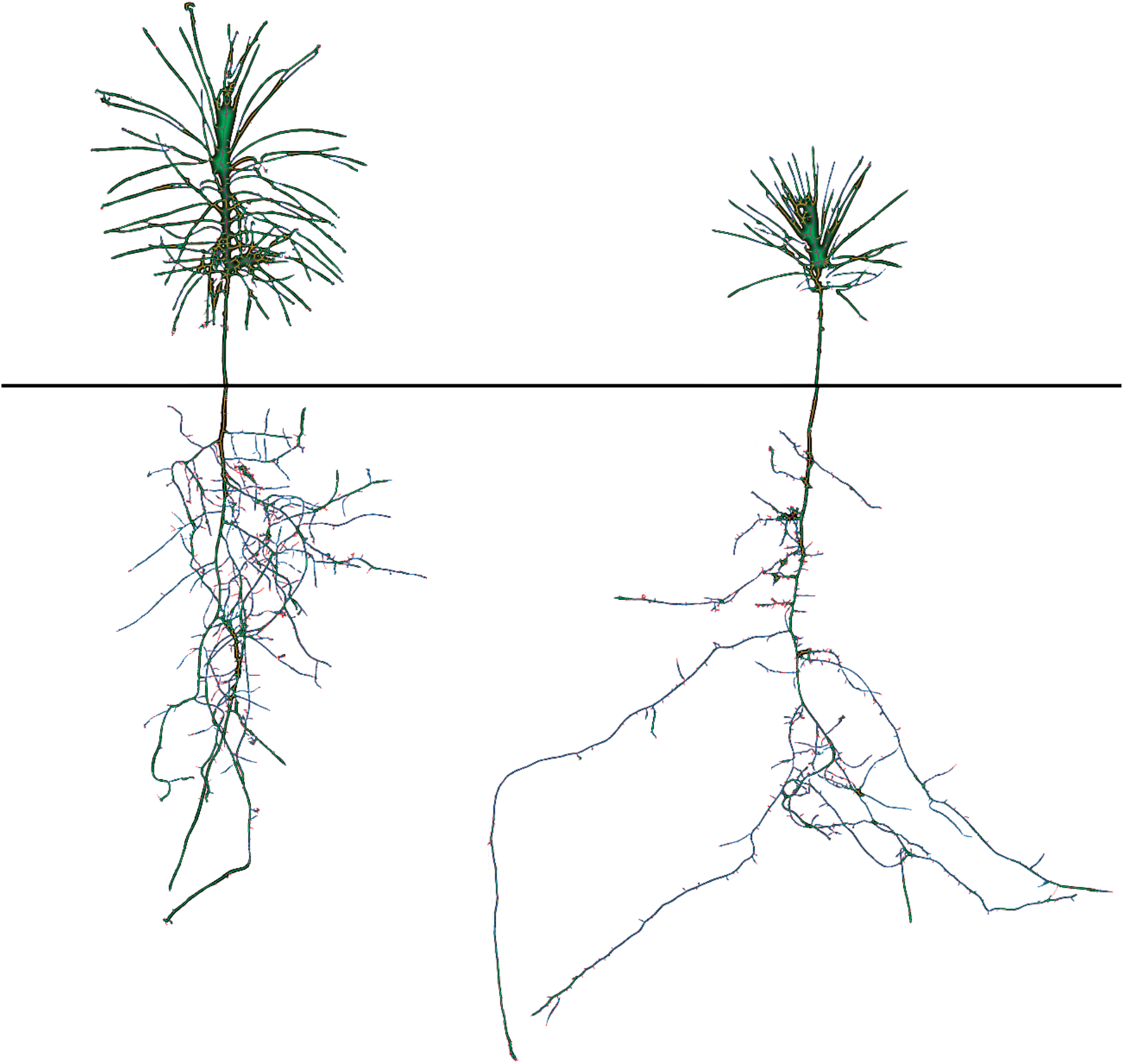
Average phenotype of plants in control (left) and drought (right) conditions. Images are segmentations of both canopies and roots of seedlings.

To further investigate the effect of drought on the root system, we represented all the root-related traits in a PCA projection for every seedling (Figure 2), both under control and drought conditions. The PC1 gradient appeared to be primarily caused by a gradient in root system density, which ranged from denser (negative values) to less dense (positive values). Meanwhile PC2 revealed a change in the root system structure from an inverse pyramid-like distribution (negative values) to a pyramid-like one (positive values). According to the projection, drought response affects root shape mainly through the reduction of root density, but also changing the spatial distribution of the upper parts. In other words, the root system under control conditions seemed to have a denser number of superficial roots.

**Figure 2.**
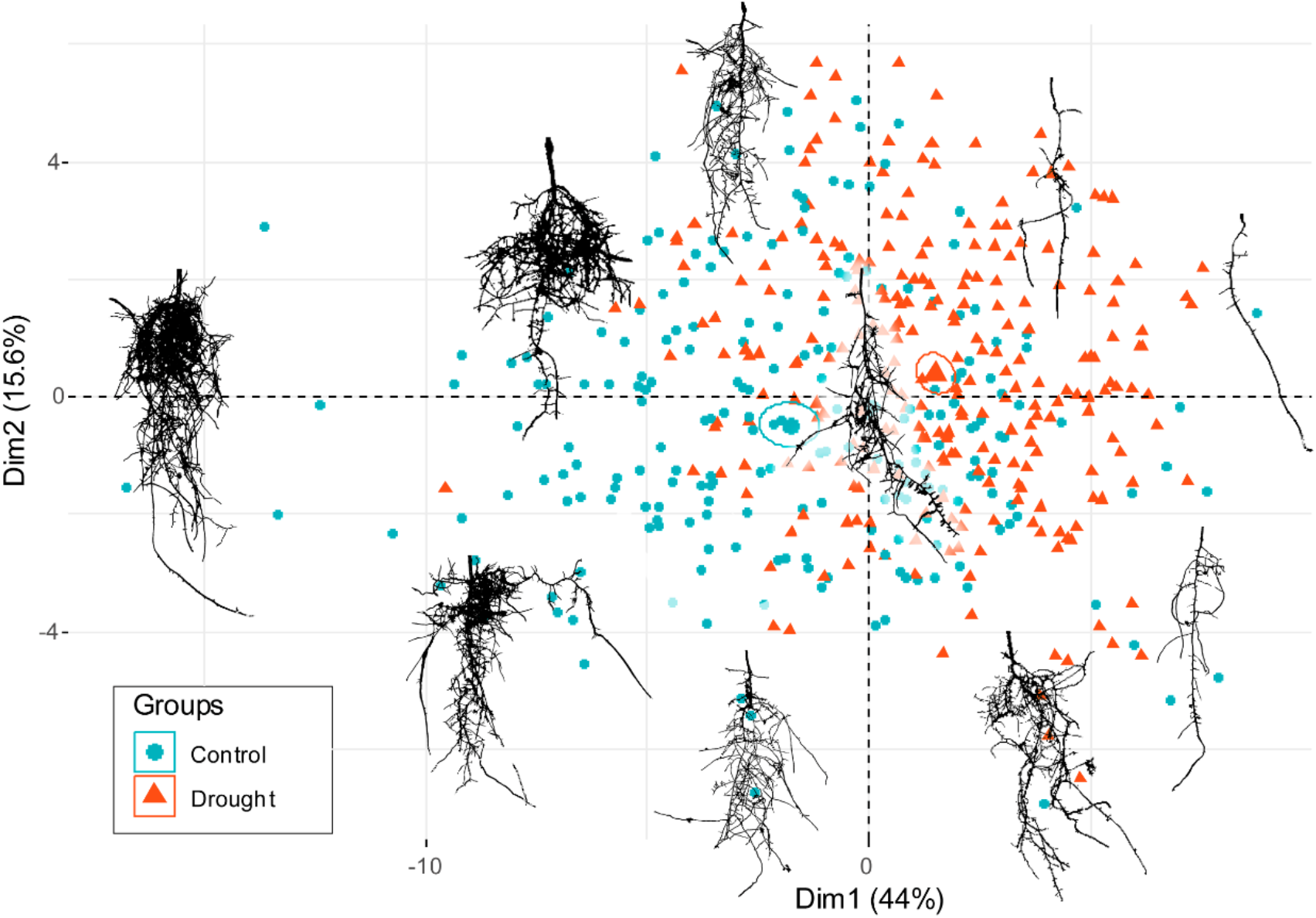
Principal Component Analysis of root parameters in Control (blue) and Drought (red) conditions for each seedling. Root scans included as reference in the image correspond to the PCA coordinates for each image.

### Defining seedling drought tolerance

To define the drought index, we used a regression analysis using the total biomass volume (i.e., canopy and roots) obtained by scanning the plants. The analysis was performed by comparing the total biomass volume per day between drought and control for all the plants belonging to the same family. The general model for the entire populations indicated that in drought conditions, an average Scots pine seedling grows 0.28*Growth in control + 3.6 mm^3^ per day (Figure 3). Family deviation from the linear model was used as drought tolerance index. In other words, the families which grew more than expected were considered tolerant, and vice versa.

**Figure 3.**
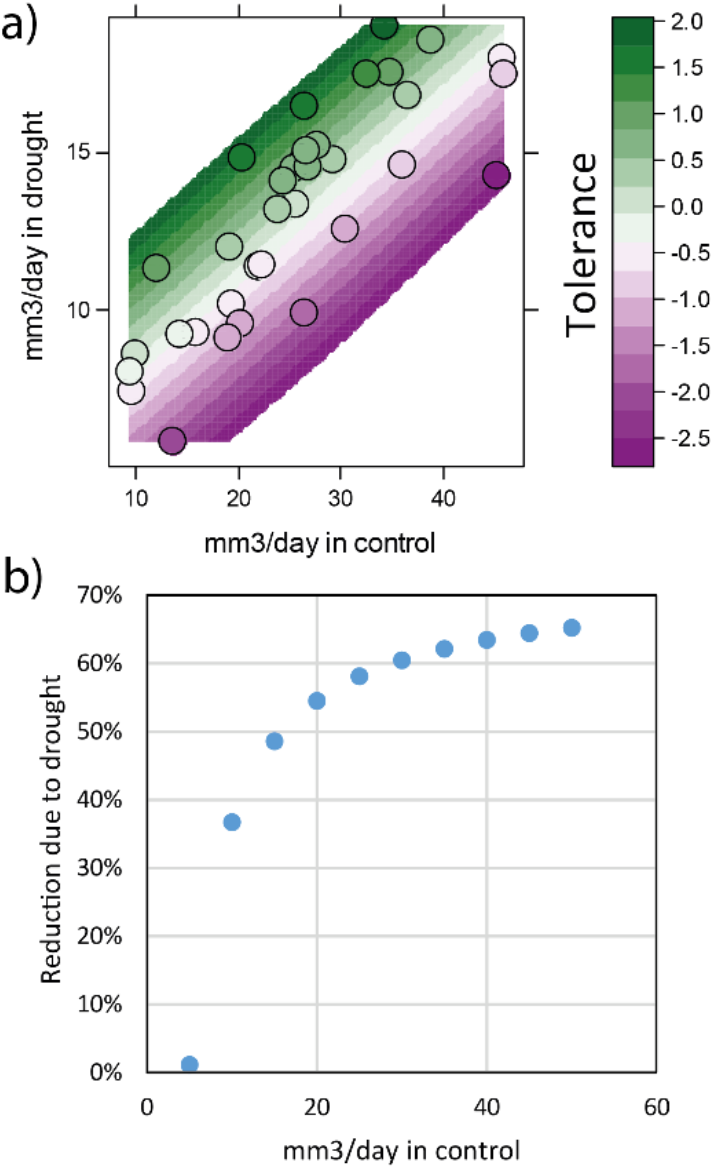
a) Relationship between the total volume in control and drought conditions in 2 month old seedlings of different families of Scots pine. Tolerance index is calculated as deviation from the linear model. Each dot is the average measurement of a family in each condition (between 4 to 12 replicates per condition). b) Simulation of the reduction of growth due to the drought effect according to the linear model versus the average growth rate of a family in Control conditions

After calculating the family average of root shape parameters PCA components by family in control and drought conditions, tolerance index parameter was included using a 2-dimensional spline interpolation (Figure 4). Under control conditions, the analysis suggests that families with lower distributed roots (smaller PC2 values) exhibited higher tolerance index when exposed to drought (Figure 4a). In other words, families with higher drought tolerance could already be selected even when grown under control conditions. The research proposes two types of ideal root shapes for drought tolerance under drought conditions: low-density roots with increased root number at the top part of the root system and a denser root system with roots mainly grouped at the bottom of the root system (Figure 4b). Regarding root reshaping response, we calculated the changes between PC and repeated the 2-d spline interpolation in order to elucidate the optimal response in root reshaping of a seedling to drought (Figure 4c). The data suggests that plants that reduce root density as little as possible and increase the proportion of roots in the top part of the system tend to have a higher drought tolerance.

**Figure 4.**
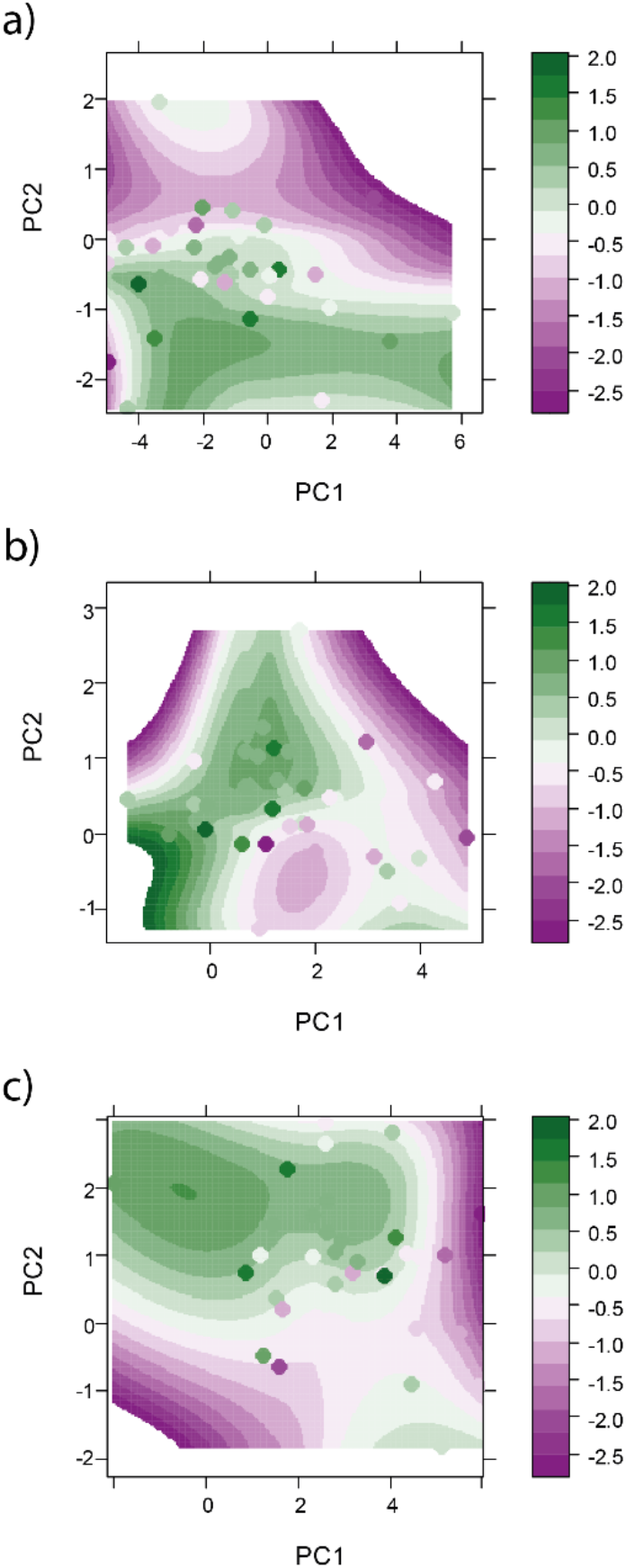
Plot of the average of Principal Components for root shape in each family and a spline interpolation of its tolerance index in control (a) and drought (b) conditions. (c) Includes the average family root shape change as difference between drought and control conditions. Background and dot color represents the tolerance index . Each dot is the average measurement of a family in each condition (between 4 to 12 replicates per condition).

Furthermore, there was some level of correlation of the family-based tolerance index with most of the individual phenotypic variables measured under control conditions (Supplementary Figure 2). Canopy Shape, Canopy Width To Depth Ratio and Median Needle Diameter, under control conditions, were significantly correlated with the tolerance parameter, but with weaker value (r^2^<0.4). On the other hand, under drought most of the parameters were positively correlated significantly with the tolerance index (Supplementary Figure 3). Amongst the parameters with a negative correlation under drought, the strongest correlation was found for Root Hole Size and Root Orientation Angle. This indicates that families with higher tolerance indexes tended to have more dense roots with smaller angles (pointing to the ground); whereas, families with a high proportion in both volume and surface of roots with diameters between 0.25 and 1 mm (ranges 2 and 3) showed a smaller drought tolerance.

### Effect of breeding on tree resilience to drought

For the traits correlated with the drought tolerance index, we observed that natural stands had lower Canopy Height, Canopy Volume, Canopy Surface Area, Canopy Perimeter, Needle Median Diameter and Needle Average Diameter in both conditions. Only in drought conditions, seedlings from natural stands exhibited lower Root Perimeter, Root Volume, Number of Root Tips, Total Root Length, Root Network Area, Root Surface Area and Root Holes in comparison to the seedligs from progeny trials (Figure 5). Under drought or control conditions, all the variables that showed a significant moderate correlation with drought tolerance showed statistical differences between natural and progeny trial stands.

**Figure 5.**
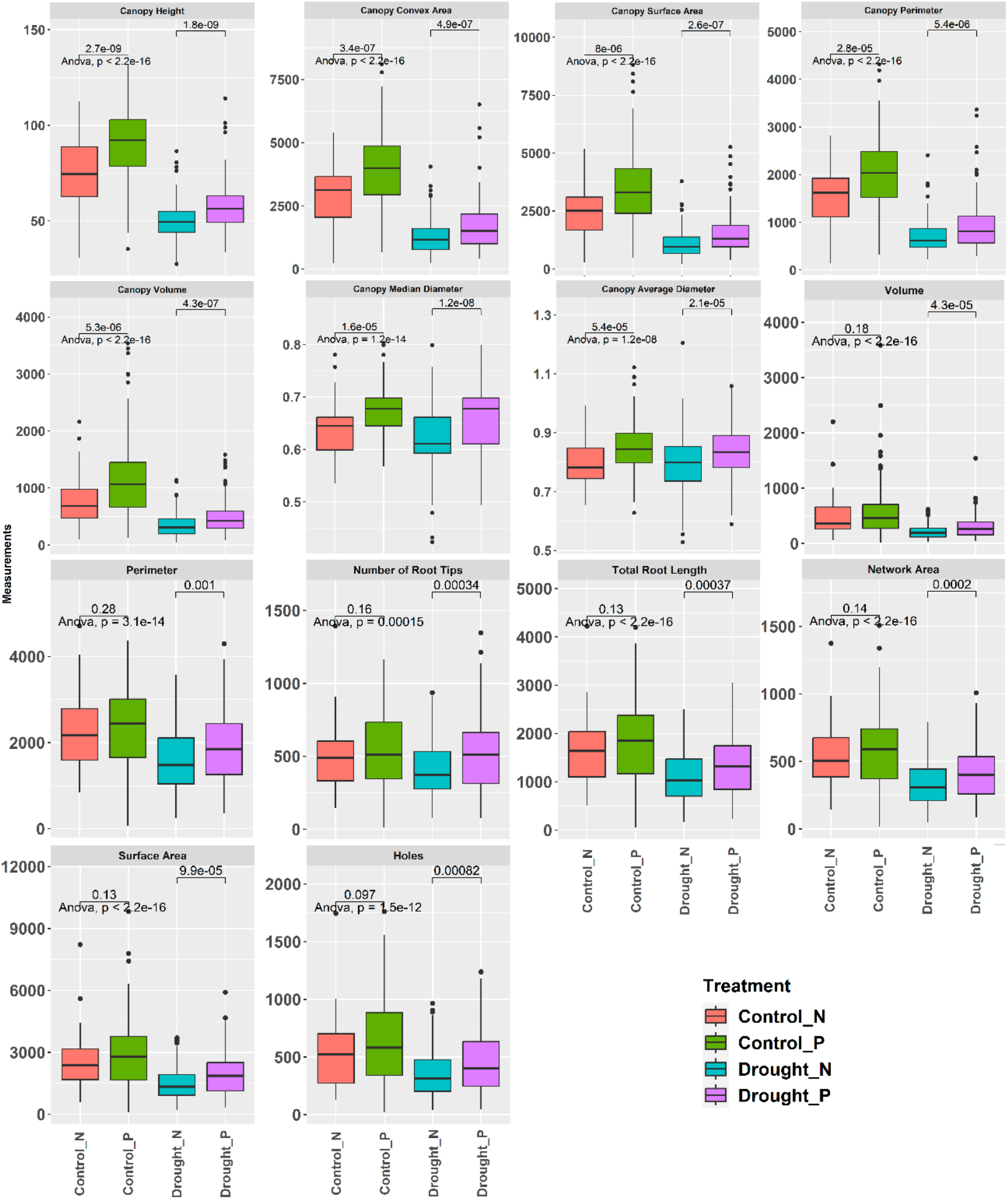
Canopy and root related trait parameters moderately correlated (r2>^i^0.4) with drought tolerance either in control or crought conditions. Statistical differences between families from natural or progeny trial stands are shown both in control and crought conditions.

A linear regression model approach was used to further study effect of selection on the response of forest trees to the drought (Figure 6). Stands of natural origin had an average reduction of growth of 24,5% (1.7 mm3/day less from 7 mm3/day baseline) in drought conditions compared to progeny trial populations indicating a significant effect of breeding (p<0.05). No interaction effect (different slope) was observed, therefore the proportional reduction of growth in drought vs control conditions is the same in all populations. Natural populations showed less average drought tolerance index than the progeny trials.

**Figure 6.**
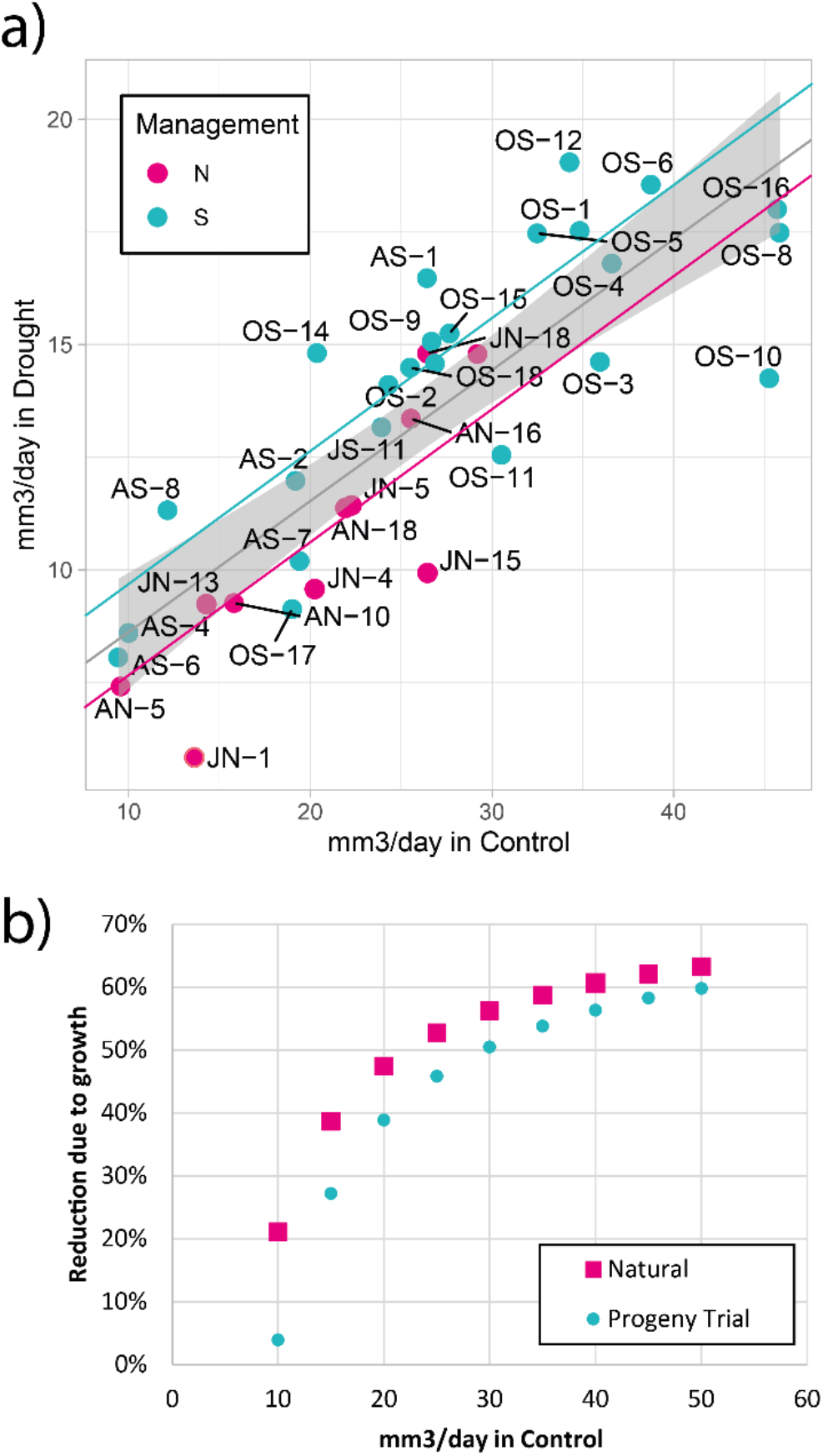
a) Relationship between total volume (canopy and root) in control and drought conditions by family. Linear model included wit 95% confidence for general average. Progeny trial stand model (in blue, S letters) and natural forest stand model (in magenta, N letters) are included. b) Simulation of the reduction of growth due to the drought effect according to the linear model versus the average growth rate of a family in control conditions for both stand origins.

### Effect of breeding on genetic diversity for drought tolerance

We estimated heritability as a measure of genetic variation for each of the measured parameter under control and drought conditions (Figure 7, Supplementary Table 3). We found that heritabilities were in general lower under drought conditions. Canopy Convex Area and Needle Median Diameter decreased the most dramatically, dropping from roughly 0.9 to 0.35. (Supplementary Table 3). On the other hand, the heritability of the Proportion of Root Surface with Diameter than 1.5mm increased under drought conditions from 0.12 to 0.33 (Supplementary Table 3).

**Figure 7.**
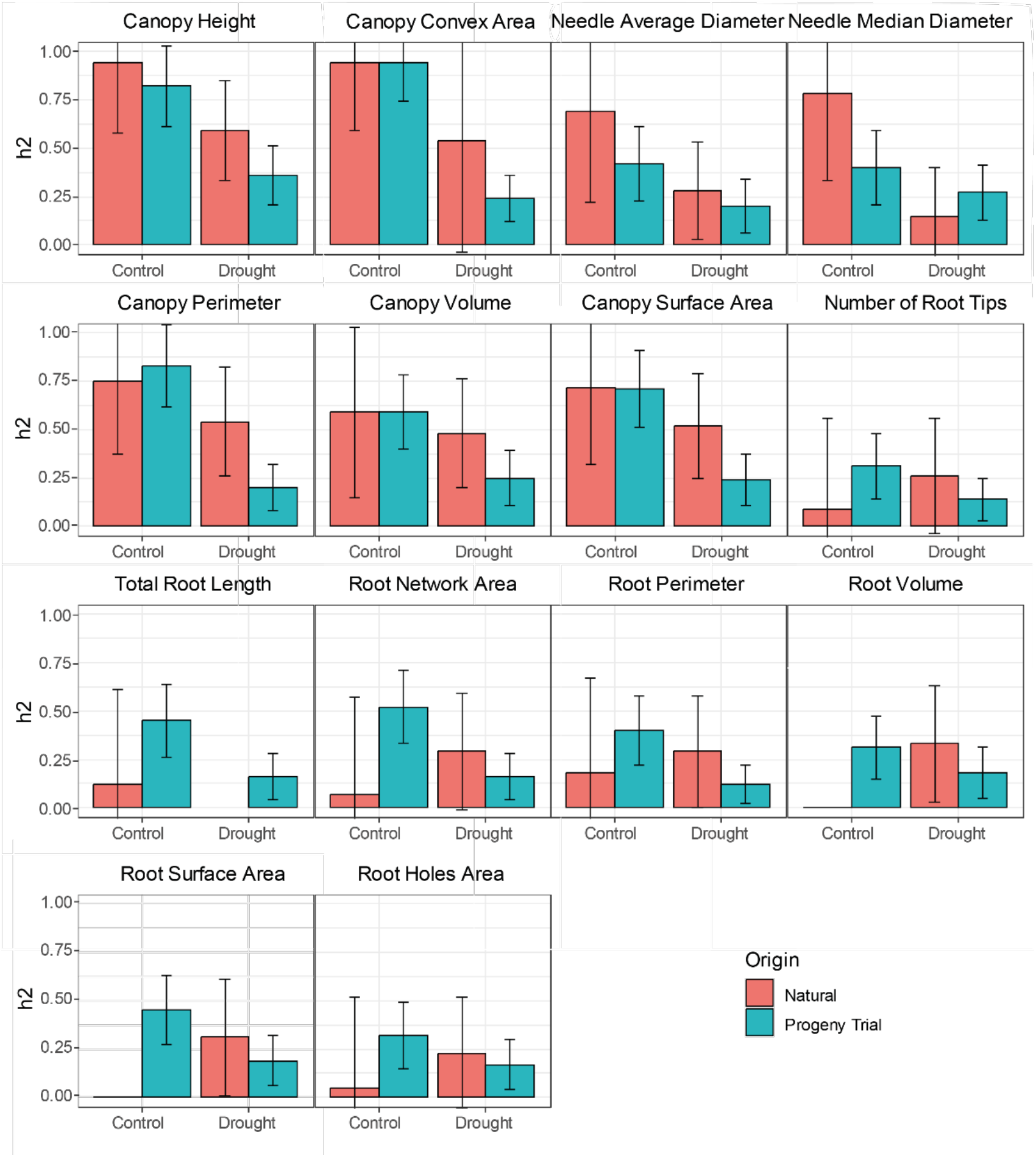
Narrow-sense heritability estimates (h^2^) of for progeny trials and natural stands tested in drought and controlled conditions.

Most of the measured parameters showed moderate to high narrow-sense heritability estimates in both natural and progeny stands, particularly for traits associated to canopy characteristics (Supplementary Table 3). In reference to the canopy parameters, the Canopy Convex Area and the Needle Median Diameter showed the highest heritability values (0.93 and 0.89, respectively) under control conditions. On the other hand, Canopy Solidity and Needle Orientation showed low heritability values. Canopy to Root Ratio, both as volume and surface, showed low heritability. Root traits had lower heritability than canopy traits, where the Total Root System Length, Network Area, Root Surface Area, Perimeter or Proportion of Roots with Diameter up to 0.5 mm showed the highest, values (more than 0.3) in control conditions. The multi-trait calculated PC1 had a moderate heritability value higher than 0.3, whereas some easily visually evaluated traits such as Root Depth, Root Solidity (density) and Width to Depth Ratio showed very low heritabilities (<0.1) (Supplementary Table 3).

For canopy traits, natural stands exhibited in general higher heritability both under control and drought conditions than the progeny stands, indicating that natural stands carry higher genetic diversity. On the other hand, regarding root traits that tendency was not as clear, instead some traits showed higher heritability in the progeny stands, such as root volume, Total Root Length or Surface Area under control conditions. Furthermore, the differences between heritabilities on Drought vs Control were less consistent than in the canopy traits. Root depth in control conditions from Natural origin showed the highest heritability value of all the traits measured. On the contrary, this trait showed very little heritability in families from progeny trials.

## Discussion

Drought in Northern Europe is projected to increase as a result of climate change, particularly during the summer (Kjellström *et al*., 2018; Spinoni *et al*., 2018). However, plant response to drought stress is difficult to assess and generalize based on existing research since the experimental designs have been carried out in a variety of ways, and no broad consensus on a single optimal methodology exists (Munns *et al*., 2010). The experimental designs described in the literature vary greatly in their design and complexity which may result in variety of possible biases due to inappropriate application of the drought treatment (Marchin *et al*., 2020). In our study, we implemented a water restriction matrix as proposed by Marchin et al. (2020) to simulate drought-restricted availability in Scots pine seedlings by sinking the water table which is the closest to the water dynamic under drought in nature.

The use of images for phenotyping complex traits was proposed already in 1970s for root shape (Rowse and Phillips, 1974). Today, a two-step approach is taken to address the overall problem of plant image-based phenotyping. The first is segmentation, which is the separation of background pixels from those belonging to the root and canopy. The second is modeling, which involves examining the relationship between those pixels to digitally rebuild the phenotypic. The first one is usually the most challenging due to its high specificity between species and conditions (Narisetti *et al*., 2022). Different segmentation algorithms can be performed using generic image-analysis software such as ImageJ or specific software for canopy traits: PlantCV, EasyPCC, PhenotyperCV, HPGA; or roots: EZ-Rhizo, SmartRoot, RootNav, ARIA, DIRT and RhizoVision Explorer. However, in many cases, currently developed software is focused on rhizotron use or intended for specific organisms (such as model plant *Arabidopsis thaliana*) architecture or organs (crown or root). Therefore, appropriate software may be a limiting factor for studies conducted under non-standard conditions and non-model organisms. Image analysis can be time-consuming, and there is often a trade-off between coarsely analyzing large datasets and conducting detailed analysis of small datasets (Lobet and Draye, 2013). Machine learning techniques are becoming a powerful resource, especially in image analysis. This technology can be applied to plant phenotyping and provide a better starting point for the previous segmentation methods that may be less accurate and more time-consuming (Ubbens and Stavness, 2017). In our case, the pipeline followed has been proven successful for analyzing both canopy and root traits in non-model species such as Scots pine.

Our study indicates that the reduction in biomass volume production rate is linear between drought and control conditions across stands and therefore the use of a deviation from this expected linear relationship can be useful for drought tolerance indicator in a controlled conditions test as previously described (Bidinger *et al*., 1982). Furthermore, a linear relationship between needle diameter and drought tolerance both in control and drought conditions could be implemented as an early selection method for drought tolerance. While this approach could become an easy-to-apply early selection method, further studies should be conducted to confirm the reliability of the method in more families and stands’ origins. Previously, other works have investigated the relationship between needle phenotype and drought tolerance. For example, needle lifespan in the tree has been previously identified as driving factor of drought tolerance in gymnosperms (Song *et al*., 2022) and changes in its morphology due to drought stress have been described and linked to hydraulic properties (Grill *et al*., 2004; Gebauer *et al*., 2015). For some traits, non-linear relationships (optimum) can obscure linear-based models. A clear example of this is the relationship between transpiration and assimilation, where there is an optimum of transpiration at the maximum assimilation rate (Brendel, 2021). According to earlier research, a non-linear models can be the best for better predicting some properties of plants under drought stress (van der Tol, Dolman, *et al*., 2008; van der Tol, Meesters, *et al*., 2008). In agreement, our multi-trait root PCA suggests that, especially in drought conditions, it exists an optimal root shape as well as an optimal root reshape response for drought tolerance rather than a purely linear correlation. Few studies have been done previously regarding genetic influence of root shape in gymnosperms. Differences have been reported regarding root length and branching in a latitudinal gradient of Norway spruce (Salmela, 2021). In Scots pine, fine roots anatomy differs according to the latitude origin of the provenances (Zadworny *et al*., 2016, 2017). Regarding drought tolerance and root architecture, in maize, a good root system to perform in drought conditions should be “Steep, Cheap and Deep” (Lynch, 2013). Similarly, our study points out that the most tolerant families have either “Steep, Cheap and Shallow” roots or either “Steep, Expensive and Deep”. Apart from water and nutrient uptake, roots have an important structural role related to abiotic stress susceptibility. Anchorage capacity is a key trait to the susceptibility of a tree to uprooting by wind damage. This is determined by the root system shape and its interaction with the soil (Dupuy, Fourcaud and Stokes, 2005). As previously mentioned, drought stress increases the susceptibility of forests to uprooting (Gardiner *et al*., 2013; Csilléry *et al*., 2017). Both rooting depth and width play a major role in the needed wind force to uproot trees (Peltola *et al*., 2011). In *Pinus pinaster*, previous studies have also highlighted the role of root shape, especially in sandy podzols (Danjon, Fourcaud and Bert, 2005) such as the ones mostly found in Scandinavia (Sauer *et al*., 2007).

Previous studies on the heritability of Scots pine features associated to needles have found values between 0.30 to 0.88 (Donnelly *et al*., 2016). However, additional gymnosperm species were shown to have low heritabilities for root-related features as root length (depth) and root:shoot ratio (Galeano and Thomas, 2023). In our study we found canopy traits to have high heritability values and moderate for root traits. Furthermore, some of the root multi-trait PCA components showed moderate heritability. Similar evidence has been presented for the genetic control of a root partitioning coefficient in *P. pinaster* (Wu and Yeh, 1997). Our study’s findings on the moderate to high heritability of traits related to drought tolerance point to the potential for early selection of drought-tolerant genotypes. This strategy is supported by the observation that conifer species’ ability to withstand drought appears to depend more heavily on young plants than on mature ones (Andivia *et al*., 2020).

One of the key objectives in tree improvement programs is to increase biomass production per breeding cycle. Our study, corroborates the positive effect of breeding on canopy and root system biomass production at the seedling stage. Interestingly, such effect on biomass seems to have resulted in an increase in the level of drought tolerance in progeny trial stands. Differences in drought tolerance between breeding generations have been previously reported (Espinoza et al., 2016; Nuhu, 2022). On the first study, a provenance-dependent relationship was found between breeding generation and drought tolerance (either positive or negative). On the second one, natural and second-generation breeding families were reported to have higher tolerance to drought than third generation breeding families of coastal Douglas-fir (Nuhu, 2022). Furthermore, the same study found that plants with good growth in control conditions tended to have better ability to overcome drought. Both this and our study differ from what has been observed in other conifers where slow growth does not ensure drought tolerance (Csilléry, Buchmann and Fady, 2020). Other studies, however, did not find any correlation between growth and drought tolerance in coastal Douglas-fir (Anekonda *et al*., 2011). These studies reflect that the effect of breeding on drought tolerance is dependent on the provenance and species and possible the selection methods. Therefore, this relationship should be studied for each specific case.

In several species, forest management through breeding has been studied for its potential source of genetic erosion (Young and Merriam, 1994; Finkeldey and Ziehe, 2004; Baucom, Estill and Cruzan, 2005; Ruņgis *et al*., 2019; Cortés, Restrepo-Montoya and Bedoya-Canas, 2020; Olsson *et al*., 2023) which is highly depends on the type of management, population size, and mating system (Ratnam *et al*., 2014). In our case, we found a decrease in genetic diversity in the natural stands for several canopy and some root traits. This consequence of intensive selection was already observed in previous studies that highlighted the potential consequences of shortterm aims in long breeding cycle species (Namkoong, 1984). Our findings support that natural forests might act as reservoirs of genetic variation. The loss in the level of genetic variation could be even higher in advanced generation breeding programs, when the main criterion of selection will be entirely growth. This effect was previously reported in coastal Douglas-fir, where differences in drought tolerance were found between second and third breeding generations (Nuhu, 2022). The increased frequency of extreme climatic events can exceed the genetic adaptability of native populations (Aitken and Bemmels, 2016). High mortality due to extreme climatic events, combined with regeneration failure, will result in the extinction of local forest populations and genetic resources (Šijačić-Nikolić, Milovanović and Nonić, 2014). An adaptive climate-smart forest management can have a central role in sustaining forests in the future (Yousefpour *et al*., 2017). Here we present evidence about the possibility of breeding different traits that can lead to a better adaptation of Scots pine to environmental stresses and also how the forest management can affect breeding potential and adaptation to present and future climatic events.

## Acknowledgements

This work was partly funded by the SLU Forest Damage Centre at the Swedish University of Agricultural Sciences (SLU) and Kempe foundation (Kempestiftelserna).

The authors acknowledge the facilities and technical assistance of the Umeå Plant Science Centre (UPSC), the Knut and Alice Wallenberg Foundation, and the Swedish Governmental Agency for Innovation Systems (VINNOVA) for their support.

